# “The Intestine is a Major Contributor to Circulating TCA Cycle Intermediates in Mice”

**DOI:** 10.1101/2022.01.20.477123

**Authors:** Wenxin Tong, Sarah A. Hannou, You Wang, Inna Astapova, Ashot Sargsyan, Ruby Monn, Venkataramana Thiriveedi, Diana Li, Jessica R. McCann, John F. Rawls, Jatin Roper, Guo-fang Zhang, Mark A. Herman

**Author notes:** **Corresponding Author:** Mark A. Herman; Duke University, 300 N. Duke Street, Durham, N.C. 27701. School of Basic Medicine, Jining Medical University, Shandong Province, China (You Wang).

## Abstract

The tricarboxylic acid (TCA) cycle is the epicenter of cellular aerobic metabolism. TCA cycle intermediates facilitate energy production and provide anabolic precursors, but also function as intra- and extracellular metabolic signals regulating pleiotropic biological processes. Despite the importance of circulating TCA cycle metabolites as signaling molecules, the source of circulating TCA cycle intermediates remains uncertain. We observe that in mice, the concentration of TCA cycle intermediates in the portal blood exceeds that in tail blood indicating that the gut is a major contributor to circulating TCA cycle metabolites. With a focus on succinate as a representative of TCA cycle intermediate with signaling activities and using a combination of germ-free mice and isotopomer tracing, we demonstrate that intestinal microbiota are not major contributors to circulating TCA cycle metabolites. Moreover, we demonstrate that the endogenous succinate production is markedly higher than intestinal succinate absorption in normal physiological conditions. Altogether, these results indicate that endogenous succinate production within the intestinal tissue is a major physiological source of circulating succinate. These results provide a foundation for investigation into the role of intestine in regulating circulating TCA cycle metabolites and related signaling effects in health and disease.

## Introduction

Animals rely on aerobic metabolism to produce chemical energy for energy intensive physiological functions like locomotion and neuronal activity. Aerobic metabolism occurs in mitochondria where acetyl-CoA, the two-carbon metabolite derived from the catabolism of carbohydrates, proteins, or lipids, is oxidized to carbon dioxide and water producing ATP via the tricarboxylic acid cycle (TCA cycle or Krebs cycle). The TCA cycle is the epicenter of cellular aerobic metabolism and TCA cycle intermediates play essential roles in this process to facilitate cellular and organismal energy homeostasis.

While the role of TCA cycle intermediates in facilitating energy production is well defined, TCA cycle metabolites can serve other roles outside of the mitochondria. For example, TCA cycle metabolites including citrate and oxaloacetate can be exported from the mitochondria and used in synthesis of glucose and lipids (1). Additionally, the TCA cycle metabolites succinate and alpha-ketoglutarate function as natural ligands for cell surface G-protein coupled receptors (GPCRs) indicating that TCA cycle derived metabolites may serve as metabolic signals (2). Succinate signals through the succinate receptor 1 (Sucnr1), a GPCR that regulates pleiotropic biological activities including urine filtration (3), immune responses (4,5), thermogenesis (6), and liver damage and fibrosis (7,8). This succinate-Sucnr1 signaling system suggests that circulating TCA cycle intermediates have the potential to regulate interorgan communication in coordination of fuel and immune homeostasis.

Although TCA cycle intermediates are produced within the mitochondria of all cells, TCA cycle intermediates are readily detected in circulation and the major sources of circulating TCA cycle intermediates in physiological conditions are less clear. Some, but not all studies have suggested that the gut microbiota is a source of circulating TCA cycle intermediates (9,10). A study in humans with obesity and type 2 diabetes observed that circulating succinate levels are positively associated with enrichment for specific succinate-generating microbial strains and suggested that gut microbiota contribute to circulating succinate (9). In contrast, colonization of mice with succinate producing bacterial species increased succinate levels in cecal matter, but did not increase portal succinate levels (10).

These studies raise questions regarding the ability of the intestine to absorb microbiome-derived and/or dietary succinate. This is of importance as chronic supplementation of succinate in the diet or drinking water impacts metabolic homeostasis in animal models, but no evidence has been provided that such supplementation studies impact circulating succinate levels (6,10). Thus, it remains unclear whether the diet or microbiota are significant contributors to circulating levels of succinate and other TCA cycle metabolites.

In this study, with a focus on succinate as a representative circulating and signaling TCA cycle intermediate, we demonstrate that endogenous gut production is a major source of circulating TCA cycle intermediates and far exceeds the dietary contribution in normal physiological circumstances. Furthermore, using both *in vivo* and *in vitro* models, we demonstrate that intestinal tissue, and not microbiota, are the predominant sources of gut-derived succinate. These results advance our knowledge of the source of circulating TCA cycle metabolites, interorgan metabolite communication, and will also provide guidance in investigating the physiological and/or pathophysiological functions of circulating TCA metabolites in health and disease.

## Methods

### Materials

Sodium succinate, cysteine, norvaline and disodium succinate-2,2,3,3-D4 (D4-succinate) were purchased from Sigma-Aldrich (14170, 30089, 53721, and 293075 respectively). D-Fructose was purchased from VWR international (VWRV0226). U-C13-fructose was purchased from Cambridge Isotope Labs (CLM-1553-PK). Dulbecco’s Modified Eagle Medium (DMEM), Advanced DMEM/F12, and PBS were purchased from Thermo Fisher (11965118, 12634010, and 10010049 respectively). Matrigel was purchased from Corning (356231).

### Animals

Animal studies were conducted according to protocols approved by Duke University’s Institutional Animal Care and Use Committee (A147-19-07 and A204-18-08). Wild-type conventionally reared C3H/HeJ and C57BL/6N animals were purchased from Jackson lab (000659, and 005304 respectively). All conventionally reared animals were fed a standard chow diet (LabDiet 5053, irradiated). Conventionally reared animals were maintained in Duke’s specific pathogen free environment with corn cob bedding. Germ-free mice were obtained from the Duke Gnotobiotic Core. These mice were originally obtained from Taconic (C57BL/6NTac). All germ-free animals were fed autoclaved Envigo 2020SX diet. Germ-free animals were maintained in a flexible film isolator (Class Biologically Clean, Madison, WI) with Alpha Dri bedding. All animals were maintained at constant temperature (~ 22°C) on a 12-hour light-dark cycle (8 AM to 8 PM). Within experiments, mice were divided randomly into different treatment groups. All experimental animals used in these studies were male, wild-type C3H/HeJ mice except for the mice used in the germ-free experiments and their controls as noted above. Except where otherwise noted with respect to specific experiments, specified treatments or euthanasia were performed at 2 PM after 5 hours food removal (9 AM to 2 PM).

### Gas chromatography-mass spectrometry (GC/MS)

Blood samples: Plasma was prepared from collected blood by centrifugation at 2,000 g, 4°C for 15 minutes. 0.1 mmol norvaline or 0.2 mM 2-Deoxy-D-glucose (2DG) was added to 10 μL plasma samples as an internal standard for the relative quantification of TCA cycle intermediates or fructose respectively. 0.5 nmol D4-succinate was added to 10 μL plasma samples as an internal standard for the absolute quantification of succinate. To 10 μL plasma samples, 400 μL MeOH, 400 μL ddH2O, and 400 uL chloroform were added sequentially and briefly vortexed after each addition. After centrifugation at 12,000 g for 15 minutes, supernatants were collected and dried under nitrogen gas.

Culture media: Culture media samples from intestinal organoids and cecal/fecal contents following fructose treatment were obtained and centrifuged at 12,000 g to remove cell debris. 0.5 mmol norvaline was added to 100 μL media samples as an internal standard for the analysis of TCA cycle intermediates. 400 μL MeOH, 400 μL ddH2O, and 400 μL chloroform were added sequentially to the samples and briefly vortexed after each addition. After centrifugation at 12,000 g for 15 minutes, supernatants were collected and dried under nitrogen gas.

Tissue and food samples: Tissue samples, and food samples were powdered under liquid N2. 400 μL ice-cold 1:1 MeOH and ddH2O were added to 10 mg powdered samples and 0.1 mmol norvaline was added in methanol as an internal standard. These samples were homogenized (TissueLyser) for 2 minutes at 30 H and centrifuged at 12,000 g, 4°C for 20 minutes. 50 μL supernatants were combined with another 350 μL ice-cold 1:1 MeOH and ddH2O prior to adding 400 μL chloroform per sample and vortexed for 30 seconds. Samples were centrifuged at 12,000 g for 15 minutes and supernatants were collected and dried completely under nitrogen evaporator.

Relative quantification and enrichment measurements of TCA cycle intermediates: Dried residues were derivatized with methoxylamine hydrochloride and tertbutyldimetheylchloros (TBDMS) sequentially as previously described [3]. Specifically, 25 μL of methoxylamine hydrochloride (2% (w/v) in pyridine) was added to the dried residues and incubated for 90 minutes at 40°C before the addition of 35 μL of TBDMS and incubation for 30 minutes at 60°C. The samples were then centrifuged for 2 minutes at 12,000 g and the supernatants of derivatized samples were transferred to GC vials for further analysis. GC/MS analysis was conducted as previously described using an Agilent 7890B GC system and Agilent 5977A Mass Spectrometer [3]. Specifically, 1 μL of the derivatized sample was injected into the GC column. GC temperature gradient started at 80°C for 2 minutes, increased to 280°C at the speed of 7°C per minute, and held at 280°C for until the completion of a run time of 40 minutes. Ionization was conducted by electron impact (EI) at 70 eV with Helium flow at 1 mL/min. Temperatures of the source, the MS quadropole, the interface, and the inlet were maintained at 230°C, 150°C, 280°C, and 250°C respectively. Mass spectra were recorded in mass scan mode from m/z 50 to 700.

Absolute quantification of succinate: Dried sample residues were derivatized with methoxylamine hydrochloride and trimethylsilyl (TMS) sequentially. Specifically, 25 μL of methoxylamine hydrochloride (2% (w/v) in pyridine) was added to the dried residues and incubated for 90 minutes at 40°C before the addition of 35 μL of TMS and incubation for 30 minutes at 60°C. The samples were then centrifuged for 2 minutes at 12,000 g and the supernatants of derivatized samples were transferred to GC vials for further analysis. GC/MS analysis was conducted using Agilent 7890B GC system and Agilent 5977A Mass Spectrometer. To be specific, 1 ul of the derivatized sample was injected into the GC column. GC temperature gradient start at 80°C for 2 minutes, increased to 280°C at the speed of 7°C per minute, and held at 280°C for until the completion of a run time of 40 minutes. Ionization was conducted EI at 70 eV with Helium flow at 1 mL/min. Temperatures of the source, the MS quadropole, the interface, and the inlet were maintained at 230°C, 150°C, 280°C, and 250°C respectively. Mass spectra were recorded in mass scan mode from m/z 50 to 700.

Relative quantification and enrichment measurements of fructose: the dried residues were derivatized with methoxylamine hydrochloride and propionic anhydride sequentially adapted from a previously described method (11). 25 μL of methoxylamine hydrochloride (2% (w/v) in pyridine) was added to the dried residues and incubated for 60 minutes at 70°C. After centrifugation for 2 minutes at 12,000 g, 50 μL propionic anhydride was added and incubated for 30 minutes at 60°C prior to another round of drying under nitrogen. The dried residues were resuspended with 55ul pure ethyl acetate and transferred to GC vials for analysis. GC/MS analysis was conducted using the Agilent 7890B GC system and Agilent 5977A Mass Spectrometer. Specifically, 1 μL of the derivatized sample was injected into the GC column. GC temperature gradient started at 90°C, increased to 260°C at the speed of 9°C per minute and further increased to 290 °C at the speed of 30°C per minute. Then temperature was held at 290°C for 5 minutes with a run time of 24.9 minutes. Ionization was conducted by EI at 70 eV with Helium flow at 1 mL/min. Temperatures of the source, the MS quad, the interface, and the inlet were maintained at 230°C, 150°C, 280°C, and 250°C, respectively. Mass spectra were recorded in mass scan mode from m/z 50 to 700.

### Liquid chromatography with tandem mass spectrometry (LC-MS/MS) for quantification of butyrate

Plasma samples were prepared the same way as described above. A modified LC-MS/MS method was developed to analyze the absolute quantity of butyrate as previously reported (12). Specifically, 30 μl plasma samples were mixed with 30 μl 20 μM 2,2,3,3,4,4,4-2H7-butyrate (D7-butyrate, internal standard) before adding to 1 ml acetonitrile to remove protein by precipitation. After centrifuge for 20 minutes at 1,200 g, supernatant was collected and transferred to a new Eppendorf tube, then dried completely by nitrogen gas. The dried residue was resuspended in a mixture solution consists of 50 μl HPLC water, 20 μl 3-Nitrophenylhydrazine hydrochloride (EDC, 120 mM), and 20 μl (N-(3-Dimethylaminopropyl)-N’-ethylcarbodiimide (3-NPH, 200 mM) for derivatization at 40 °C for 30 minutes. After centrifuge for 10 minutes at 1,200 g in room temperature, supernatant was transferred to an LC-MS/MS vial for butyrate analysis. LC-MS/MS was conducted using a Sciex QTRAP 6500+ MS connected with a Sciex AD UHPLC. Specifically, an Agilent C18 column (Pursuit XRs C18 150 × 2.0 mm, 5 μm) was employed to generate a flow rate at 0.4 ml/min for separation. Gradient methods were conducted within two mobile phases. Mobile phase A was composed of 98% H2O and 2% acetonitrile with 0.1% formic acid. On the other hand, mobile phase B was consisting of 98% acetonitrile and 2% H2O containing 0.1% formic acid. At first, gradient started with 98% phase A and 2% phase B for 30 seconds. After 7.5 minutes, phase B was increased to 90% while phase A was decreased to 10% at 8 minutes. After maintaining at 90% phase B and 10% phase A for another 4.5 minutes, phase B was returned to initial 2% within 0.5 minutes. Finally, the column was re-equilibrated for 9 minutes with 98% phase A and 2% phase B before the next injection. The injection volume was 3 μl. MRM in negative mode was used for butyrate assay. The MS/MS parameters were set as following: curtain gas: 35 psi, source temperature: 600 °C, Gas 1: 55 psi, Gas 2: 55 psi, CAD: 10, Ion spray voltage: −4500 V, EP: −10 V, and CXP: −14.

### Stable Isotope Analysis

Isotopologues that containing 0, 1, to n heavy atom(s) in a molecule were referred as M+0, M+1, to M+n respectively. The retention times of individual metabolites and their isotopomers are described in Supplementary Table 1. Isotope enrichment and labeling analysis in this study were corrected for natural isotope distribution as previously described (13,14). Specifically, the stable isotope enrichment of each metabolite in plasma, tissue, organoids, and fecal/cecal content was corrected based on the natural isotope distribution measured in the same type of samples treated without labeled molecule sources. For example, the natural isotope distribution matrix of individually measured metabolite in the blood samples was experimentally determined by assaying and averaging the blood samples from animals who had not received any treatments involving isotopically labeled sources. Similarly, the natural isotope distribution matrix of individually measured metabolites in the organoids samples was determined by assaying and averaging the blood samples from organoids that were incubated with PBS without exposure to isotopically labeled sources.

### Intestinal Organoid Culture

Intestinal crypts were isolated from 8-week old male mice and used to grow enterocyte-enriched intestinal organoids as previously reported using ENR-growth media [Advanced DMEM/F12 supplemented with 50 ng/mL murine EGF, 1 μM N-acetylcysteine, 100 nM GSK269962, 1XB27, 2% Noggin conditional media, and 2% R-spondin-1 conditional media] (15). Briefly, small intestines were removed, incised longitudinally, and cut into 2- to 5-mm fragments before being washed with ice-cold PBS 3 times. Washed intestinal fragments were then incubated with ice-cold PBS containing 10 mM EDTA for 30 minutes on a rocker at 4 °C. Tissue was removed from solution containing EDTA and placed in ice-cold PBS and mechanically shaken by hand for 30 seconds. Crypts were enriched by filtering through a 70-uM cell strainer to remove the villi. Enriched crypts were resuspended in ENR-growth media after centrifugation at 300 g for 5 minutes. After manually counting, crypts were further diluted and embedded in Matrigel (3:1) and seeded on 12-well plates at a density of approximately 900 crypts per well and incubated with ENR-growth media. After 5 days of proliferation and differentiation, intestinal organoids were treated with either 10 mM glucose or 10 mM U-C13-fructose for overnight (5 PM to 9AM) at 37 °C. Immediately after incubation with glucose or fructose, the media were harvested for measurement of succinate labeling using GC/MS as described above.

### Fecal/Cecal Content Culture

Fecal and cecal contents were freshly harvested from the anus and cecum respectively from 8-week-old male mice and incubated with or without 10 mM U-C13 fructose in 0.05% cysteine supplemented PBS for 30 minutes at 37 °C. Immediately after incubation, culture media from the fecal and cecal content samples were prepared for further GC/MS analysis as described above.

### Statistics

All data are presented as mean ± SEM. Analysis of TCA levels between the tail and portal blood were conducted using a paired student’s T-test in GraphPad Prism. Comparisons of metabolite levels between time points following oral bolus gavage were analyzed using 1-way ANOVA and Tukey’s test. As appropriate, data were analyzed for statistical significance with GraphPad Prism using 2-way ANOVA and individual comparisons with a Tukey’s test. Statistical significance was assumed at P less than 0.05.

## Results

### The gut contributes to circulating TCA cycle metabolites

The TCA cycle and its intermediates are essential for ATP production in mitochondria under aerobic conditions. As meal-derived nutrients are typically absorbed from the intestine and delivered to the liver prior to mixing with systemic circulation, we sampled blood from the portal vein and tail vein of mice to establish the baseline levels of the TCA cycle intermediates. We observed that relative levels of succinate, fumarate, and malate are ~ 9-fold higher in the portal blood when compared to tail blood (Figure 1A). Citrate levels were also higher in portal compared to tail blood, but only ~ 2-fold higher (Figure 1A). Conducting absolute quantification of succinate in the portal versus tail plasma using a labeled succinate internal standard confirmed the increase in portal blood demonstrating a succinate level of 150.4 ± 20.9 μM in portal blood compared to 51.9 ± 17.2 μM in tail blood (P < 0.001, Figure 1B). This marked portal-peripheral gradient indicates that circulating TCA cycle intermediates are gut-derived, and either absorbed from the intestinal lumen or generated from the intestinal tissue.

**Figure 1.**
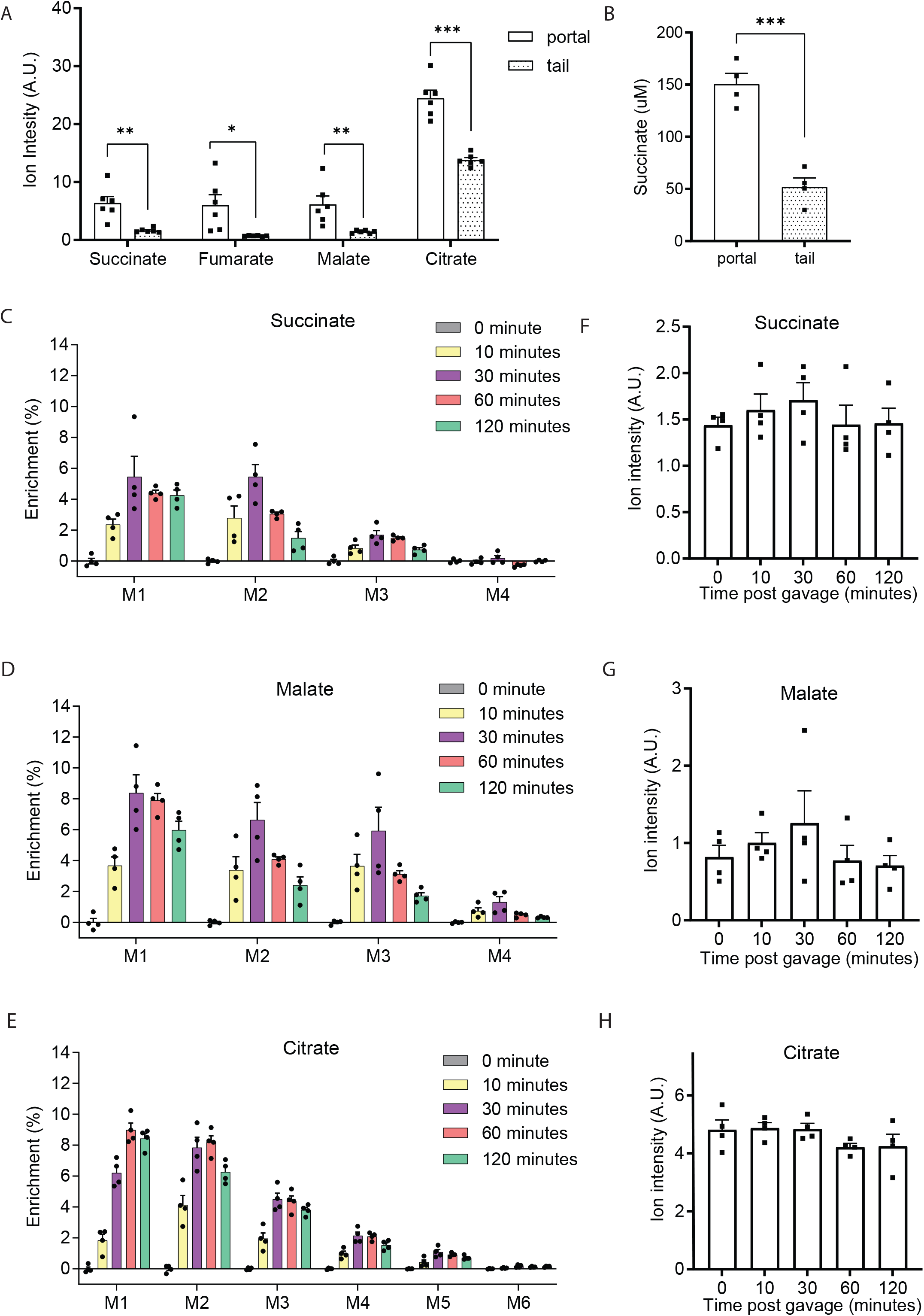
The gut is a contributor to circulating TCA cycle intermediates. Relative quantification of TCA cycle metabolites (A) and absolute quantification of succinate (B) in portal versus tail plasma samples from 8-weeks old, male mice after 5 hours food removal. Norvaline and D4-sodium succinate were used as internal standards for relative and absolute quantification measurement by GC/MS, respectively (n=6 per group). Enrichment (C, D, E) and relative quantification (F, G, H) of portal TCA cycle intermediates after oral-gavage in 8-weeks old male mice with U-C13-fructose (0.48 g/Kg body weight, n=4 per group). Data represent means ± SEM. * p<0.05; ** p<0.01; *** p<0.001. Analysis in A and B performed via paired student t-test. Analysis in panel F – H performed via one-way ANOVA and Fisher’s LSD for post-hoc analysis comparing to baseline (0 minutes).

To examine the contribution of gut carbohydrate metabolism to portal TCA cycle intermediates we orally gavaged mice with U-C13-fructose (0.48 g/kg) as fructose can be metabolized at high rates by both microbiota and enterocytes in the gut (16,17). The relative quantification and enrichment of TCA cycle intermediates in the portal blood were assessed by GC/MS at intervals for 2 hours following gavage. The rapid appearance of labeled carbons in portal TCA cycle intermediates supports the hypothesis that orally administered fructose can be readily metabolized into TCA cycle intermediates in gut (Figure 1C, D, E). While fructose-derived carbons were readily incorporated into portal TCA cycle intermediates, the quantity of the TCA cycle intermediates (succinate, malate, and citrate) in the portal vein remained relatively constant over this two-hour period (Figure 1F, G, H). The increase in isotope enrichment without changes in TCA cycle intermediate levels indicate that TCA cycle intermediates are constitutively secreted from the gut into circulation although the source of substrate for TCA cycle intermediate production is variable.

### Microbiota are not essential to maintain portal succinate

TCA cycle intermediates found in portal blood appear to be constitutively produced in the gut. The cellular origin of these TCA cycle intermediates is uncertain as both the intestinal tissue and microbiota residing in the gut have the capacity to produce them. Given interest in succinate as a metabolic signal (18), we focused on succinate as a representative TCA cycle intermediate in the remainder of this study. We measured succinate levels in blood samples of conventionally reared mice to age- and strain-matched mice that were reared in a germ-free environment. Butyrate is a well-established circulating metabolite produced largely by microbiota (19). As expected, portal butyrate levels are 143-fold higher in the portal blood of the conventionally reared mice compared to the germ-free mice (Figure 2A). In contrast, succinate levels in the portal blood of germ-free mice were similar to that of conventionally reared animals (Figure 2B). Thus, intestinal microbiota are not a major source of circulating succinate.

**Figure 2.**
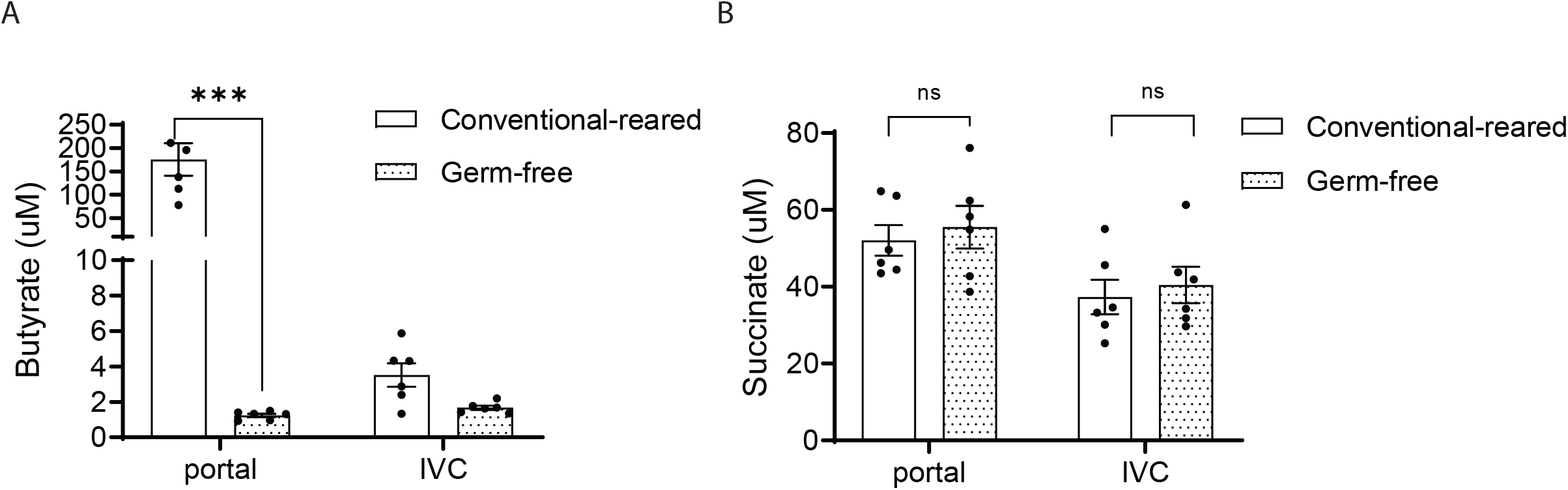
Intestinal microbiota are not a major source of circulating succinate. Absolute quantification of butyrate (A) and succinate (B) in plasma from the portal vein tail versus inferior vena cava of 12-week old germ-free male mice versus their conventionally reared mice. Butyrate levels were measured by LC-MS/MS using D7-butyrate as internal standard. Succinate levels were measured by GC/MS using D4-succinate as internal standard (n=6 per group). Data represent means ± SEM. * p<0.05; ** p<0.01; *** p<0.001. Data were analyzed by two-way ANOVA and Bonferroni analysis for post hoc comparisons.

### Labeled precursors produce distinct succinate labeling patterns when exposed to intestinal organoids versus microbiota

Although intestinal microbiota are not essential for intestinal succinate production, this does not exclude the possibility that microbiota can contribute to circulating succinate in conventionally bred animals, and that other undefined mechanisms maintain portal succinate levels. Whereas eukaryotes generate succinate from carbohydrate precursors exclusively via the TCA cycle, microbiota could generate succinate from carbohydrate precursors via the glyoxylate shunt independent from the TCA cycle (20) (Figure 3A). Thus, the labeling pattern of succinate produced from C-13-labeled carbohydrate-derived three-carbon substrates are likely to be different when produced in mammalian cells versus microbiota (20). Specifically, mammalian cells using predominantly the TCA cycle will generate M+2 succinate in the first turn of the cycle and M+1 succinate in the second turn of the cycle, leading to predominant enrichment in M+1 and M+2 succinate (Figure 3A). In contrast, microbiota use a highly active PEP carboxykinase which converts PEP into oxaloacetate and then condenses acetyl-CoA and oxaloacetate via isocitrate lyase into isocitrate which is used to generate glyoxylate and succinate (20). The succinate generated in this pathway from C-13-labeled carbohydrate-derived three-carbon substrates will be enriched at M+3 and M+4 in addition to M+1 and M+2 (Figure 3A).

**Figure 3.**
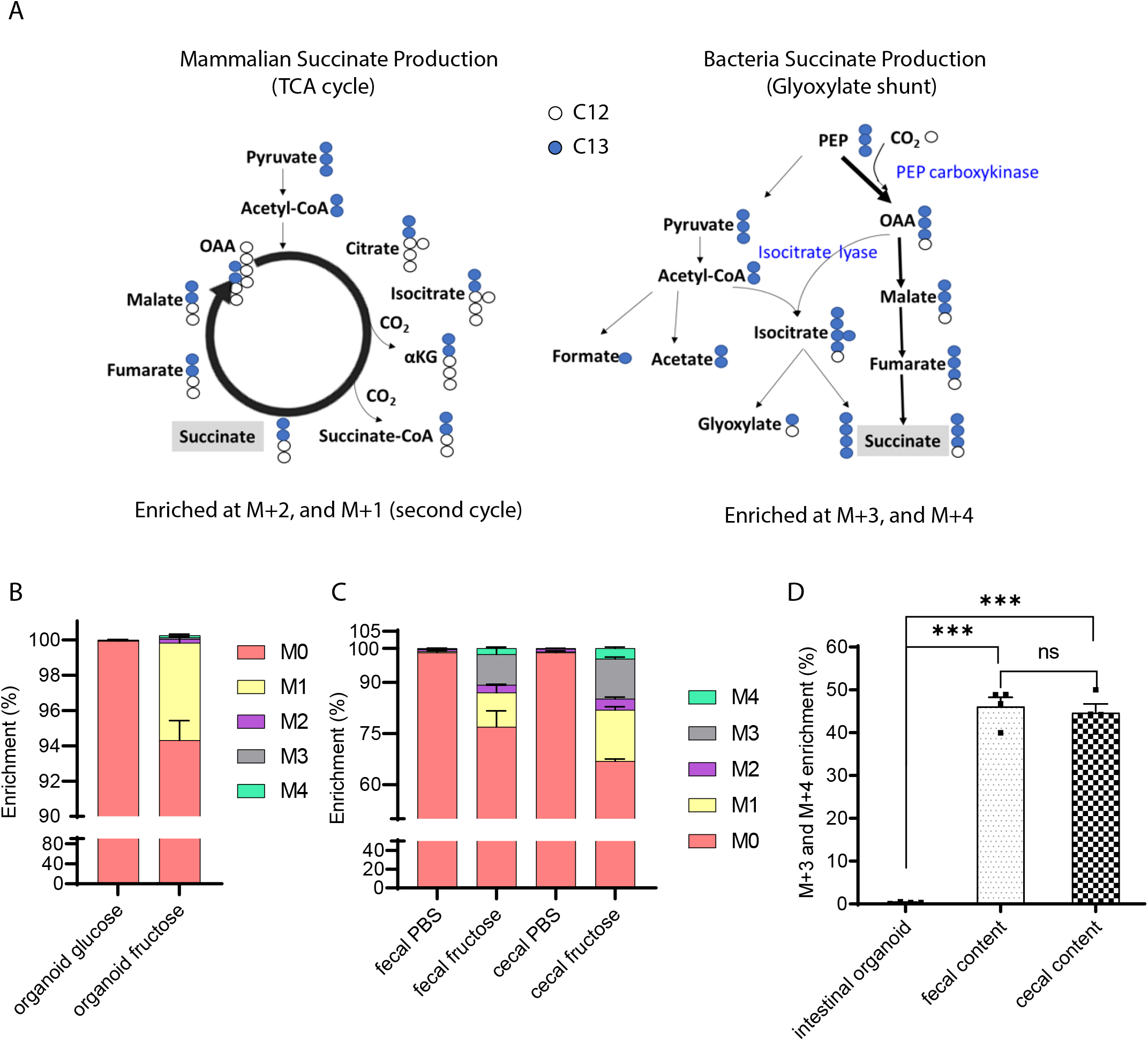
Intestinal organoids compared with cecal and fecal contents generate distinct succinate labeling patterns from labeled precursors. (A) Theoretical isotope-labeling patterns of succinate produced from C13-labeled three carbon substrates in mammalian cells versus bacteria. Measured labeling pattern in (B) mammalian cells and (C) microbial enriched samples. Intestinal crypts and fresh cecal and fecal contents were harvested from 8-weeks-old male mice. Intestinal organoids were differentiated into enterocytes prior to treatment with 100 mM U-C13-fructose for 24 hours (n=3 per group). Freshly harvested cecal and fecal contents were cultured with 100 mM U-C13-fructose for 30 minutes (n=4 per group). (D) Quantification of the ratio of M+3 and M+4 succinate to total labeled succinate in cultured intestinal organoids and cecal/fecal contents described in (B) and (C). Data represent means ± SEM. * p<0.05; ** p<0.01; *** p<0.001. Data were analyzed by one-way ANOVA and Fisher’s LSD for post-hoc analysis between intestinal organoids, fecal contents, and cecal contents.

We examined succinate enrichment within fecal and cecal contents in vitro and in cultured intestinal organoids after incubation with U-C13-fructose. As expected, isotope-labeled succinate was identified in both the intestinal organoids and fecal and cecal contents when cultured with U-C13-fructose. In the intestinal organoid experiment, the M+1 and M+2 succinate accounts for 83.7% of the labeled succinate (Figure 3 B), consistent with the hypothesis that mammalian cells predominantly use the TCA cycle to generate succinate from carbohydrate precursors. In contrast, fecal and cecal contents generated a much higher proportion of M+3 and M+4 labeled succinate, accounting for nearly 44.8% to 46.6% of the labeled succinate (Figure 3 C-D). These substantial differences indicate that the contribution of microbiota versus mammalian tissue to succinate production can be distinguished based on labeling patterns following administration of labeled carbohydrate precursors.

### Portal succinate is produced from intestinal tissue and not the microbiota

We next compared the succinate labeling pattern in portal blood, intestinal tissue, and cecal contents in mice after providing large amounts of labeled precursors in the form of U-C13-fructose. The fructose dose, 4 g/kg, is higher than the amount that mice can fully absorb in the small intestine, thus providing adequate labeling in both intestinal tissue and the bulk of intestinal bacteria present in the cecum and colon (16,21). Portal blood, intestinal tissue, and cecal contents were harvested from animals that were not gavaged or at either 10 minutes or 2 hours after gavage. The two time points were chosen based on the liquid transition speed in typical mice. Specifically, 10 minutes is sufficient for liquid to transition to the small intestine, but not cecum, whereas 2 hours is aimed at ensuring cecal fructose metabolism (16,21). The portal succinate labeling pattern and specifically the proportion of M+1 and M+2 labeled succinate consistent with production in the eukaryotic TCA cycle were similar in portal blood and intestinal tissue, and distinct from the pattern in the cecal contents at both time points (Figure 4 A-B). These data demonstrate that portal succinate is predominantly produced by intestinal tissue and not microbiota.

**Figure 4.**
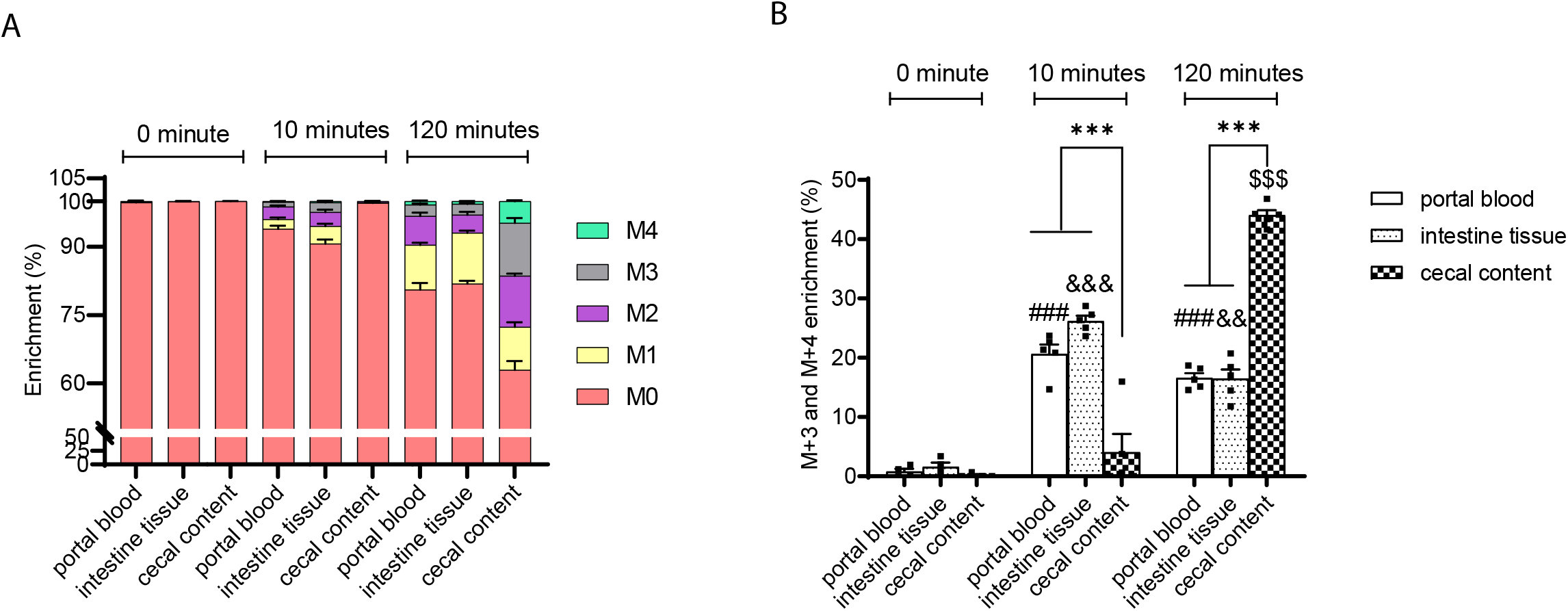
The succinate labelling pattern in portal plasma is similar to that in intestinal tissue while distinct from that in the cecal contents following gavage with U-C13-fructose. (A) The enrichment and (B) the ratio of M+3 plus M+4 to total labeled succinate in the portal plasma, intestinal tissue, and cecal contents after gavage of 8-weeks-old male mice with U-C13-fructose (4g/Kg body weight) (n=4 per group at each time point). Data represent means ± SEM. * p<0.05; ** p<0.01; *** p<0.001. Data were analyzed by two-way ANOVA and Tukey’s test was applied for post hoc comparisons. # portal plasma comparisons across time points, & intestinal tissue comparisons across time points, $ cecal content comparisons across time points, * comparisons between tissues within time points.

### Compared to endogenous production, intestinal succinate absorption capacity is limited

Prior studies have suggested that dietary succinate supplementation can regulate multiple systemic biological activities through the effects of absorbed succinate on the Sucnr1 receptor in diverse tissues (6,22,23). However, there is limited data demonstrating that oral or dietary succinate supplementation increases circulating succinate levels (24). Our data indicates that succinate produced by microbiota in the large intestine is not readily absorbed, but this does not exclude the possibility that dietary succinate could be absorbed in the small intestine. Thus, we sought to examine the intestinal capacity for absorption of orally administered succinate and compare this to the rate of endogenous intestinal succinate production.

Portal succinate levels are a mixture of succinate absorbed from the intestinal lumen, succinate produced endogenously within intestinal tissue, and succinate carried into the portal circulation via systemic recirculation or via production in other tissues. The content of succinate in typical human or mouse diets is not well defined. Succinate may be added to foods as a flavor enhancer at levels not to exceed 0.08% (g/g) in condiments and less than 0.006% (g/g) in meat products per FDA guidelines (21CFR184, Sec. 184.1091). By GC/MS measurement, the succinate content of a standard mouse chow diet (LabDiet 5053), is 146.5 μg/g comprising less than 0.015% of food by weight (Figure 5A).

**Figure 5.**
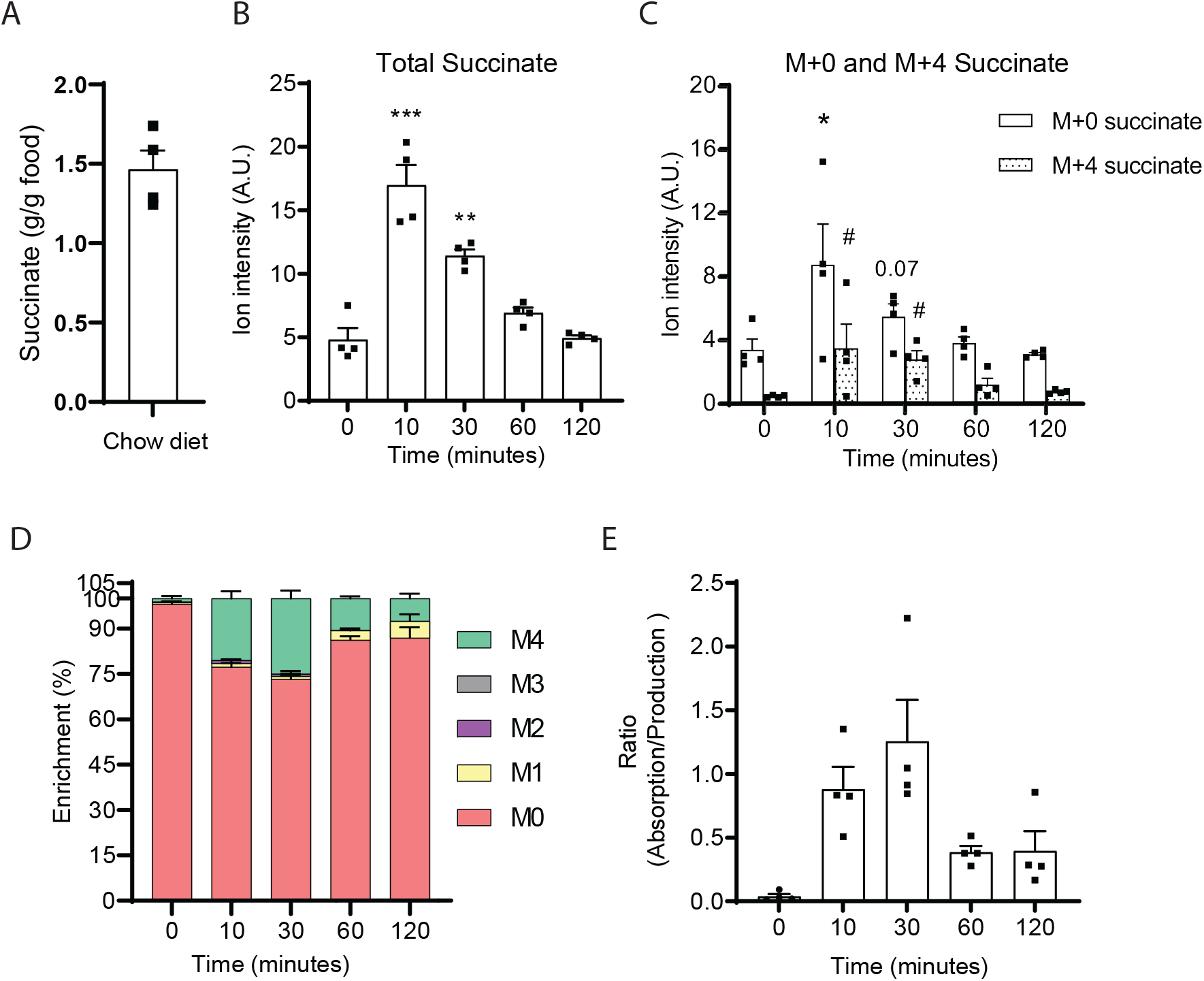
The intestinal absorption of succinate is limited. (A) Succinate content of normal chow diet (LabDiet 5053). (B) Total succinate ion counts and (C) M+0 and M+4 isotopomers ion counts and (D) enrichment in portal plasma after gavage of 1:1 D-4 succinate and unlabeled succinate (1.46 g Na Succinate/Kg body weight). Norvaline was used as internal standard. (E) The estimated ratio of intestinal absorption to endogenous production of succinate. (n=4 at each time point). Data represent means ± SEM. * or # p<0.05; ** or ## p<0.01; *** or ### p<0.001. Data were analyzed by one-way ANOVA and Fisher’s LSD for post-hoc analysis between baseline (0 minutes) and time points post D4-succinate administration; In panel C, * indicates the comparison of M+0 succinate and # indicates the comparison of M+4 succinate with the zero-time point.

To define the upper limits of intestinal succinate absorption and to compare this to the rate of endogenous gut succinate production, we gavaged mice with 1:1 disodium succinate-2,2,3,3-D4 and unlabeled disodium succinate at a dose of 1.46 g/kg which is close to the solubility of succinate in water. This amount of succinate (~ 36.5 mg for a 25g mouse) is more than 60-fold higher than the amount of succinate consumed by a mouse on a standard chow diet in a day (~ 0.6 mg succinate for a mouse that eats 4g food per day). After gavage of this supraphysiological succinate bolus, total portal succinate levels increased within 10 minutes and declined to baseline levels after ~ 1 hour (Figure 5B). This suggests that succinate can be absorbed from the intestinal lumen. Interestingly, at the time point of 10 minutes, the increase magnitude for M+0 succinate (from 3.35 ± 1.32 to 8.70 ± 5.10 A.U.) is higher than M+4 succinate (from 0.41 ± 0.08 to 3.44 ± 2.99 A.U.), suggesting the osmolarity stress of the supraphysiological succinate bolus may promote the secretion of endogenous gut-derived succinate (Figure 5C). During the 2 hours’ time period, the maximum proportion of M+4 succinate accounts for total succinate only range from 20 to 25% (Figure 5D). This indicates that the absorbed exogenous 1:1 labeled to unlabeled succinate was substantially diluted by endogenous unlabeled succinate (Figure 5E). Using our model (see Supplemental Figure 1 and associated Methods), we estimated that at peak absorption of this supraphysiological succinate bolus, the contribution of endogenous succinate and absorbed succinate are approximately equal (Figure 5D). These results indicate that in normal physiological circumstances, where succinate is present in ingested foods in limited amounts, the vast majority of circulating succinate is produced endogenously and that succinate absorption likely contributes negligibly to portal succinate levels.

## Discussion

Recent studies illustrate important roles for TCA cycle intermediates outside of their role in energy production including important roles for circulating TCA cycle intermediates as metabolic signals (2,18). This field has advanced with the discovery of cell-surface receptors sensitive to specific TCA cycle metabolites including succinate and alpha-ketoglutarate (25). While the role of metabolites like succinate in regulating cellular function and organismal homeostasis is rapidly advancing, few studies have assessed the source of circulating TCA cycle intermediates.

We observed that the levels of TCA cycle intermediates are markedly higher in the portal blood than the peripheral circulation indicating the gut is a major source of circulating TCA cycle intermediates (Figure 1A, B). Recently Jang and colleagues used steady-state stable isotopic tracer infusions to assess sites of production and consumption of a wide range of metabolites including TCA cycle metabolites in pigs (26). Based on sampling of the pancreaticoduodenal vein, they concluded that the pancreas was a major source of circulating TCA cycle metabolites. However, as this vein carries venous blood from both the pancreas and duodenum, these investigators could not discriminate whether the duodenum versus pancreas is the actual source. Moreover, in this study analysis of blood obtained from the splenic vein which drains the pancreas tail was not consistent with the pancreas as a source of TCA cycle intermediates.

Fructose is a dietary sugar that is preferentially metabolized in selected cell types including enterocytes in the intestine, hepatocytes in the liver, and proximal tubule cells in the kidney (27,28). Fructose is not readily metabolized in the pancreas (27). Gavaging mice with labeled fructose rapidly and robustly labels TCA cycle intermediates in the portal vein and to comparable degrees within duodenal tissue (Fig 1 C-D and 4 A). Although multiple intestinal epithelial cell types (enterocytes, goblet cells, and paneth cells) may express the Glut5 luminal fructose transporter and fructolytic enzymes (29), these are expressed at high levels in absorptive enterocytes which are thought to mediate the bulk of intestinal fructose metabolism (27,30). Altogether, these results indicate that enterocytes in the proximal intestine are major contributors to gut-derived circulating TCA cycle metabolites.

According to studies conducted by us and others, the physiological circulating succinate levels range from 0 to 50 μM in humans (9), and 0 to 100 μM in rodents (22). This range of succinate coincides with the pharmacological range (EC_50_) of succinate receptor (Sucnr1) activation – 20 to 50 μM in human and 10 to 100 μM in rodents (2). Thus, changes in the circulating succinate level within the physiological range have potential to affect the activity of Sucnr1 and its downstream signaling. This supports succinate’s potential role as an important circulating metabolic signal.

While our results indicate that enterocytes are major sources of circulating succinate in physiological circumstances, other tissues may also play important roles. For example, acute cellular hypoxia such as myocardial infarction, ischemia of the kidney, or liver ischemia are known to result in succinate production, and this may increase circulating succinate levels (31–33). Exercise-induced hypoxia in skeletal muscles can produce substantial amounts of succinate which can transiently increase circulating succinate levels (34). Chronic, mild increases in circulating succinate are associated with cardiometabolic diseases including obesity, hypertension, and type-2 diabetes and the source of this succinate is unclear (9,35). Additional studies will be required to determine whether intestinal TCA cycle metabolite production is a contributor to changes in circulating succinate associated with pathological conditions.

Intestinal microbiota are increasingly recognized as a major source of bioactive metabolites in systemic circulation which may have pleiotropic physiological effects (36). As examples, microbiota-derived short-chain fatty acids and acetate both play important roles in hepatic gluconeogenesis and de novo lipogenesis participating in diet-induced metabolic disease (17,37). Serena et al. observed an enrichment of succinate-generating microbiota in association with circulating succinate levels in patients with metabolic diseases and hypothesized that the microbiota contributes to this increase (9). De Vadder et al. subsequently transplanted one of these succinate-generating microbial strains, *P. copri,* into conventional mice and observed significantly increased succinate in cecal contents in association with improved glucose tolerance (10). However, this was not associated with increased circulating succinate indicating that increased levels of succinate in the intestinal lumen are not sufficient to increase circulating succinate levels (10). De Vadder et al. interpreted this data to suggest that microbially-derived succinate is absorbed and consumed within the intestine as gluconeogenic substrate. However, intestinal gluconeogenesis is largely restricted to the small intestine whereas succinate production from microbiota occurs predominantly in the colon (38). As we cannot rule out the possibility of nutrient exchange between the small and large intestine, we assessed portal succinate levels in conventional and germ-free mice (Figure 3B), directly demonstrating that gut microbiota are not essential for maintaining circulating succinate levels. Moreover, by comparing portal succinate labeling patterns after oral administration of labeled substrate, we demonstrate that intestinal tissue rather than microbiota are the predominant contributor to portal succinate in conventional mice (Figure 4 A, B). Altogether, we and others have demonstrated that although microbiota can generate substantial amounts of succinate in the intestinal lumen, this microbiota-produced succinate is distinct from that produced endogenously in the small intestine. Thus, microbiota are not a major contributor to circulating succinate. These results raise questions as to the causal nature of the improvements in glucose homeostasis after transplantation of succinate-generating microbiota. At a minimum, the improvements in metabolism associated with this transplant cannot be attributed changes in circulating succinate as a metabolic signal.

These studies also raise questions about the capacity of the intestine to absorb succinate from the lumen. This question is important as several studies have suggested that dietary supplementation of succinate can increase circulating succinate and regulate biological functions through activation of SUCNR1 in multiple organs including the liver and adipose tissue (6,22,39). However, none of these studies assessed the impact of dietary succinate supplementation on circulating succinate levels. Here, we examined intestinal succinate absorption capacity and determine the relative importance of succinate absorption compared to endogenous production.

To approach this problem, we first sought literature regarding the succinate content of typical diets and were surprised to find no clear reference for this in typical human or animal diets. Hence, we measured succinate levels a standard mouse chow diet and found that succinate contributes no more than 0.015% g/g diet. We elected to gavage mice with a quantity of labeled succinate that far exceeds the typical dietary content. At this extreme, supraphysiological dose, succinate absorption transiently approached the levels of endogenous succinate production (Figure 5 B-E). These results suggest that the modest amounts of succinate present in typical diets contributes negligibly to gut-derived circulating succinate levels.

While the results from this study support the conclusion that enterocytes are a major source of circulating TCA cycle metabolites, there are several limitations. The model used to compare the relative rates of succinate absorption versus endogenous production relies on several assumptions which include: 1) metabolism of absorbed and endogenously produced succinate follows first-order kinetics, and 2) the recirculation of labeled succinate is negligible. First, although most of metabolites follow first-order elimination kinetics (40), it is possible that succinate elimination actually follows zero-order elimination rule. However, the rapid peak and return to baseline after within 1 hour of gavage is inconsistent with a zero-order elimination model. Second, although we disregard the recirculation of intestinal absorbed labeled metabolites in our model, this assumption could lead to an overestimation of the absorption of succinate Thus, if recirculation is substantial, the actual intestinal absorption rate would be even smaller than what we estimated here and would further support our conclusion that intestinal absorption of succinate makes a minor contribution.

In summary, intestinal tissue is a major source of circulating TCA cycle metabolites. The role of intestinally-derived circulating TCA cycle intermediates as metabolic signals remains to be investigated in future studies.

## Author Contributions

**MAH, WT, SAH, RM, and IA** designed, performed, and interpreted the conventional mouse experiments. **YW and GZ** designed, performed, and interpreted the germ-free mouse experiments. **GZ, WT, MAH, YW, and SAH** designed, performed, and interpreted metabolomics analysis. **JFR, JRM, WT, and MAH** designed and performed the *in vitro* culturing of cecal/fecal content. **VT, DL, and JR** assisted with the intestinal organoid system, and **SAH, WT, and MAH** designed and performed the intestinal organoid experiments. **AS and RM** assisted with performing and interpreting experiments. **MAH** conceived of, designed, and supervised the experimental plan, and interpreted experiments. **WT** and **MAH** wrote the manuscript. All authors edited the manuscript.

## Acknowledgements

This work was supported by NIH grants 5R01 DK121710 (MAH). Metabolomics support was provided by the North Carolina Diabetes Research Center (NCDRC) (P30 DK124723).

## Notes

**Conflict of interest statement:** Mark A. Herman receives research support from Eli Lilly, and Co.

### Competing Interest Statement

Mark A. Herman receives research support from Eli Lilly, and Co.

