## Supplemental for "“The Intestine is a Major Contributor to Circulating TCA Cycle Intermediates in Mice”"

**Supplementary Table 1**

| Metabolites and their isotopomers | m/z | Retention time (min) |
|---|---|---|
| M+0 Succinate (Relative quantification and enrichment analysis-TBDMS method) | 289.1 | 18.0 |
| M+1 Succinate (Relative quantification and enrichment analysis-TBDMS method) | 290.1 | 18.0 |
| M+2 Succinate (Relative quantification and enrichment analysis-TBDMS method) | 291.1 | 18.0 |
| M+3 Succinate (Relative quantification and enrichment analysis-TBDMS method) | 292.1 | 18.0 |
| M+4 Succinate (Relative quantification and enrichment analysis-TBDMS method) | 293.1 | 18.0 |
| M+0 Succinate (Absolute quantification-TMS method) | 247.0 | 10.8 |
| M+4 Succinate (Absolute quantification-TMS method) | 251.0 | 10.7 |
| M+0 Fumarate | 287.1 | 18.4 |
| M+1 Fumarate | 288.1 | 18.4 |
| M+2 Fumarate | 289.1 | 18.4 |
| M+3 Fumarate | 290.1 | 18.4 |
| M+4 Fumarate | 291.1 | 18.4 |
| M+0 Malate | 419.3 | 23.0 |
| M+1 Malate | 420.3 | 23.0 |
| M+2 Malate | 421.3 | 23.0 |
| M+3 Malate | 422.3 | 23.0 |
| M+4 Malate | 423.3 | 23.0 |
| M+0 Citrate | 459.3 | 29.0 |
| M+1 Citrate | 460.3 | 29.0 |
| M+2 Citrate | 461.3 | 29.0 |
| M+3 Citrate | 462.3 | 29.0 |
| M+4 Citrate | 463.3 | 29.0 |
| M+5 Citrate | 464.3 | 29.0 |
| M+6 Citrate | 465.3 | 29.0 |
| Norvaline | 288.2 | 16.6 |
| M+0 Fructose | 328.0 | 19.8 |
| M+1 Fructose | 329.0 | 19.8 |
| M+2 Fructose | 330.0 | 19.8 |
| M+3 Fructose | 331.0 | 19.8 |
| M+4 Fructose | 332.0 | 19.8 |
| M+5 Fructose | 333.0 | 19.8 |
| M+6 Fructose | 334.0 | 19.8 |
| 2DG | 182.1 | 17.1 |

**Supplementary Figure 1 and Associated Methods:**

The crude relative quantification model used here is based on standard isotope dilution methodologies. Specifically, mice were gavaged with 1:1 isotope-labeled and unlabeled metabolites. Animals were then sacrificed over a 2-hour time period with the following time points: 0 (no gavage), 10, 30, 60, and 120 minutes for the harvest of portal blood. The relative quantity and labeling pattern of metabolites in the portal blood samples were measured by GC/MS.

(Supplementary Figure 1A and 1B) Metabolites in the portal blood are a mixture of metabolites transported into portal circulation following absorption as well as recirculation of metabolites from systemic circulation include metabolites derived from endogenous production which may occur in the intestine or other tissues. Thus, the labeling of absorbed isotope-labeled metabolites are diluted by the endogenously produced metabolites and recirculating metabolites which can be assessed in portal blood samples. The endogenously produced metabolites are assumed to be exclusively unlabeled because the natural isotopomer labeling of their substrates is negligible (41). One assumption made by in this model is that the redistribution of intestinal absorbed metabolites is limited during our acute oral gavage experiment. Ignoring redistribution may overestimate the absorption rate since all labeled metabolites were assumed to be from absorption. We also assumed that the metabolism of absorbed nutrients in intestinal tissues follows first-order kinetics, which is approximately true for most drugs and nutrients (40). Thus, the fraction of transported labeled metabolites following absorption will be the same as transported unlabeled metabolite from endogenous production within the intestine ($E_A$ / A = $E_P$ / P, $E_A$: Elimination of absorption; A: Absorption; $E_P$: Elimination of production; P: Production). As we gavaged a bolus of metabolites at a 1:1 labeled to unlabeled ratio, the transported labeled metabolites are proportional to half of the total absorbed metabolite ($T_{labeled} \propto ½ T_A$, $T_{labeled}$: Transported labeled metabolite). Transported unlabeled metabolites originate from both external absorption and endogenous production. Transported unlabeled metabolites are proportional to the sum of half trasnported absorption and transported intestinal production ($T_{unlabeled} \propto (½T_A + T_P)$, $T_{unlabeled}$: Transported unlabeled metabolites). Combining these equations, we crudely estimate the ratio of external absorption to endogenous production using the transported labeled and unlabeled metabolites amount ($A/P = E_A/E_P \propto (2 * E_{labeled}) / (E_{unlabeled} - E_{labeled})$). The quantity and labeling of metabolites are measured by GC/MS in portal blood samples.

We tested this model using fructose, taking advantage of the fact that fructose is quickly absorbed and readily metabolized in the intestine and that endogenous fructose production is likely small relative to fructose absorption in mammals (28). We measured (Supplementary Figure 1C) the relative quantification and (Supplementary Figure 1D) enrichment of portal fructose after oral-gavage with 1:1 U-C13-fructose and unlabeled fructose (0.48 g/Kg body weight) (n=4 per time point). As expected, within 10 minutes of fructose gavage, both portal and tail fructose levels rapidly and robustly increased approximately 50-fold and 20-fold above their baseline levels, respectively (Supplementary Figure 1C). The relative abundance of M+6 fructose was similar to that of M+0 fructose at the 10-minute time point and comparable to the 1:1 ratio of labeled and unlabeled fructose that was provided by gavage. The lack of dilution of M+6 fructose with M+0 fructose indicates that endogenously produced fructose was not a major contributor to portal fructose at this time point. The gradual decline in M+6 compared to M+0 fructose from 30 minutes on is indicative of the dilution of the gavaged mixture of labeled and unlabeled fructose with endogenously produced fructose (Supplementary Figure 1D). Based on the relative quantification model, we estimate that the rate of intestinal fructose absorption is approximately 25-fold greater than the rate of endogenously produced fructose production at the 10-minute time point (Supplementary Figure 1E). These findings are consistent known aspects of fructose metabolism in mice (27) and illustrate that this simplified relative quantification model is useful in estimating the relative amounts of absorbed versus endogenously produced substrates *in vivo*.

Data represent means ± SEM. * p<0.05; ** p<0.01; *** p<0.001. Data in panel A were analyzed by two-way ANOVA with time and location as co-variable Bonferroni analysis for post hoc comparisons; # portal comparisons between time points, & tail comparisons between time points, * comparisons between tail and portal. Data in panel C were analyzed by one-way ANOVA and Fisher's LSD for post-hoc analysis of the ratios of M+6 to M+0 fructose between baseline (0 minutes) and time points post fructose administration (10, 30, 60, 120 minutes).

41. Nilsson, R. (2020) Validity of natural isotope abundance correction for metabolic flux analysis. *Mathematical biosciences* **330**, 108481

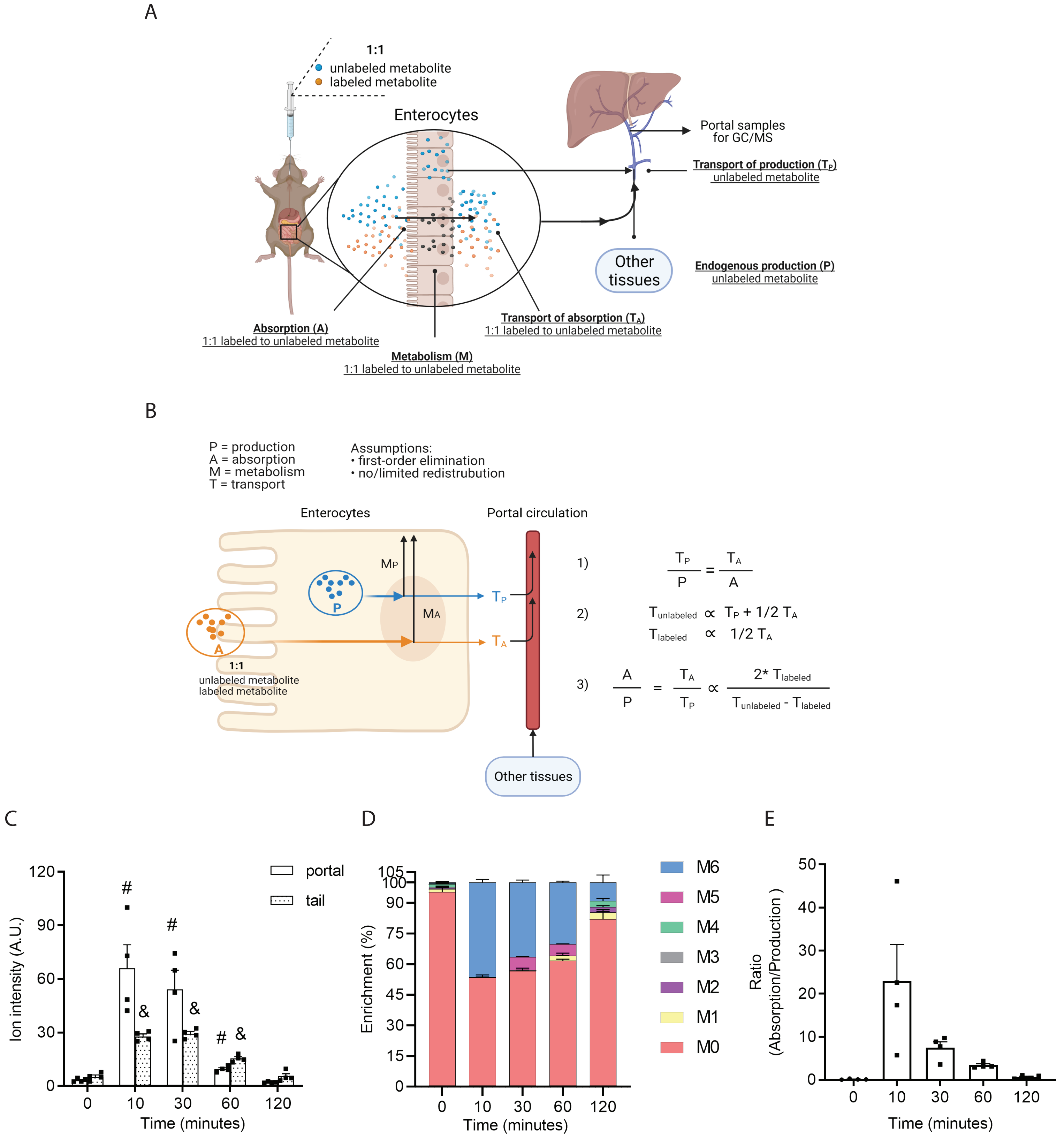


Supplementary Figure 1.
